# MeiCOfi: Meiotic CrossOver Finder in haploid, diploid, polyploid and hyper-recombinant genomes

**DOI:** 10.64898/2026.04.29.721680

**Authors:** Roven Rommel Fuentes, Joiselle B. Fernandes, Tamara Susanto, Yazhong Wang, Charles J. Underwood

## Abstract

During the meiotic cell division, homologous chromosomes pair and recombine, leading to large reciprocal exchanges of genetic information. In most species, meiotic crossovers (COs) are crucial for normal chromosome segregation and they generate genetic diversity, which can be acted upon by natural selection in wild populations or by breeders to combine desirable traits in a genome. Identifying the position and frequency of COs is therefore essential in both classical genetics studies and breeding programmes. However, a computational tool capable of accurately detecting COs across diverse contexts, including varying marker densities, genome size and structure, recombination rate, and ploidy, remains lacking. We developed *MeiCOfi* (Meiotic CrossOver Finder) to detect meiotic crossover events at high-resolution from low-coverage genome sequencing data. We evaluated it using data from *Arabidopsis thaliana*, rice, barley and both intra- and inter-specific tomato hybrids, encompassing a wide range of genome complexities and marker densities. It reliably detects crossovers in hyper-recombinant *A. thaliana* with up to 62 CO per backcross offspring and in haploid gametes from barley with sequencing coverage as low as 0.1x. It can identify crossovers in polyploid genomes, including simulated recombinant tetraploids and also real data from tetraploid tomato hybrid offspring. Our results demonstrate that *MeiCOfi* can robustly identify crossovers in diverse genomic contexts.

## Introduction

Eukaryotic species typically reproduce by sexual reproduction, which requires the reduction of genome ploidy through meiosis, such that diploid organisms produce haploid spores and gametes. During the first meiotic division, homologous chromosomes pair and synapse, and large reciprocal exchanges of genetic information occur via meiotic crossovers (COs). COs break down genetic linkage in wild populations, generating variation upon which natural selection may act, while in artificial breeding programmes, they are crucial to incremental genetic gain. Traditionally COs were identified by cytological analysis of chromosome meiotic spreads, such as by manual counting of chiasmata or recombination sites^1-4^. Apart from cytogenetic techniques, COs can be detected using genetic approaches. Recombination in a particular region was detected in seeds or pollen through fluorescent tagging^5^. Moreover, early genetic maps were generated using markers based on probes and sequence tags^6-8^. In the era of high-throughput genotyping and genomics, high-density linkage maps have been generated, for example in tomato and maize^9,10^. Genomics-based methods rely on the polymorphism between the parental genomes to infer recombination events. As the cost of genome sequencing declined, sequencing of recombinant offspring became a preferred method to detect genome-wide crossovers at higher-resolution^11-14^. To avoid the laborious growing of an offspring population, alternative approaches identified COs by sequencing pooled gametes from an F1 parent^15-17^.

Despite the central role of meiotic crossovers in heredity, evolution, and breeding, bioinformatic tools that robustly work in variable levels of genomic heterozygosity and across ploidies are lacking. Coalescent- or linkage disequilibrium-based methods have been developed to estimate population-scaled recombination rates from a natural population^18,19^. Moreover, for detecting recombination in an offspring population, one of the early-developed tools is ReCombine, which was developed for recombinant yeast^20^. TIGER was developed based on Hidden-Markov-Model to detect recombination in low-coverage resequenced genomes and was tested on *A. thaliana*^21^. Other detection methods like inGap-family^22^ and MRLR^23^ were optimized on *A. thaliana* or human genomes. The majority of these tools were developed for a particular species or ploidy but do not transfer well to highly divergent crosses or larger plant genomes. Some prioritize sensitivity in low coverage samples more than reporting high-resolution COs. In this paper, we present a method that can detect crossovers at high-resolution across species, ploidy levels and recombination rates.

## Materials and Methods

### Plant material growth and generation

Seeds of *S. lycopersicum* cv. Moneyberg-TMV (MbTMV), *S. cheesmaniae* accession LA1039, and *S. pennellii* accession LA0716 were pre-treated in 700 ppm GA_3_ (Gibberellic acid) for 20 hours (to enhance germination) and then rinsed with sterilised water. Seeds were further subjected to sterilization in saturated Na_3_PO_4_ (trisodium phosphate) for 10 minutes and rinsed thrice with sterile water. Subsequently, seeds were treated with 2.7% NaOCl (sodium hypochlorite) for 30 minutes and rinsed again 5 times. All seeds were then sown on 0.7% agar plates and incubated under a 16 h/8 h photoperiod at 26°C/22°C day/night conditions. After germination, seedlings were transferred to MS (Murashige and Skoog) medium (4.3 g/L MS salts, 100 mg/L myo-inositol, 20–30 g/L sucrose, 7 g/L PhytoAgar, pH 5.9) and grown in jars, before transfer to soil. Flowers of MbTMV were emasculated and independently pollinated by pollen collected from LA0716 and LA1039. Hybrid seeds were germinated and grown as above.

### Generation of tetraploid tomato hybrids

Diploid F1 hybrids of *Solanum lycopersicum* cv. Moneyberg-TMV and *Solanum cheesmaniae* accession LA1039 were grown *in vitro* in MS media (as above) and young leaves (3-4 weeks old) were subjected to chromosome doubling following the method of Chetelat, et al. ^24^. Regenerated shoots were screened for ploidy by flow cytometry and non-chimeric rooted tetraploids (Mb-Che tetraploid hybrids) were transferred to soil in the greenhouse and Mb-Che 4x F2 seeds were harvested. 13 F2 seeds germinated and young true leaf material was collected for DNA sequencing

### DNA Sequencing

Young true leaf material was lyophilised overnight in a freeze dryer (Martin Christ GmbH). DNA extraction was carried out on a KingFisher robot (Thermofisher) using the NucleoMag Plant Kit (Macherey and Nagel, P/O 744400.4). Transposase-based libraries (Tn5, Nextera) were prepared and sequenced on Illumina NextSeq 2000 and NovaSeq 6000 platforms. A total of 95 Mb-Pen diploid F2 plants and 13 Mb-Che tetraploid F2 plants were sequenced.

### Marker selection

*MeiCOfi* accepts as input the ratio of reads supporting each parental SNP allele. SNPs are detected by aligning the reads of one parent against the assembly of the other using *bwa-mem*^*25*^ and then running *GATK-Haplotypecaller*^*26*^ for variant detection. In the absence of assembled genomes for both parents, reads from both parents are aligned against a reference genome, and the mutually exclusive SNPs detected in both parents are selected. If sequencing data of the F1 hybrid parent is available, it is used to identify discordant allele ratios and remove problematic markers. Tracts of homopolymers in the reference genome are detected and used to remove possible false SNPs or artefacts of misaligned reads in low complexity regions. The final list of SNPs is used in *bcftools*^*27*^ to genotype samples and generate an input file for *MeiCOfi*. The alignment file can be provided as input to filter markers in regions with coverage significantly deviating from the genome average, avoiding problematic markers due to repetitive regions or false positive COs due to the change in allele frequencies in regions with copy number variations or aneuploidy.

### Identify putative crossover region

From the input read counts, allele frequency (AF) is computed per marker followed by the average AF per window to average out erroneous markers. A minimum number of markers is required per window. The average allele frequencies in two consecutive blocks of windows is compared to estimate the frequency change. The difference between the left block (AF *l*_*i*_ = [*i*-*n*+1,*i*], where *n* is the minimum number of windows) and the right block (AF *r*_*i*_ = [*i*+*1,i*+*n*]) may indicate a possible site for recombination. Depending on the ploidy of the sample, the expected frequency change is computed, such as 1.0, 0.5 and 0.25 for haploid, diploid, and tetraploid, respectively. The region in between the two consecutive blocks with frequency change (ΔAF = (*r* - *l*)*(ploidy) above a threshold will serve as a candidate CO location for the boundary refining step. The threshold may be adjusted to account for the distribution and density of SNPs in the sample. By inspecting the flanking markers, the number of crossovers in each candidate region can be estimated.

### Refining the crossover position

The raw frequencies of SNPs overlapping the candidate region are retrieved to refine the boundaries of the crossover. Using an R function for *changepoint* detection^28^, the raw SNP frequencies and the estimated number of COs, *MeiCOfi* detects in finer resolution the crossover location. Crossover location is further refined by computing the distances between the frequencies of SNPs (*m* number of SNPs per block) in two consecutive non-overlapping blocks and identifying the local distance maxima. In case a finer location cannot be identified due to the quality of AF and density of markers, the initial boundaries of the candidate crossover are reported.

### Simulating genomes

Using the genome assemblies of *Solanum lycopersicum* cv. Moneyberg-TMV (MbTMV)^29^ and *Solanum pennellii* (LA0716)^30^, paired-end data of the parental genomes were simulated using art_modern^31^. Random CO locations in the genome were selected and used to subset reads from the parental data, simulating a recombinant haplotype. Depending on the ploidy level being simulated, the recombinant haplotype was mixed with a proportional set of reads from either or both of the parents. For each coverage level (0.5x to 8x), 100 samples were simulated. Reads were aligned against the MbTMV reference genome using *bwa-mem*, followed by variant calling using bcftools.

Crossover lists from ten F2 samples (Mb-Pen diploid) were manually validated using the allele frequency plots. To simulate genomes with lower marker density, these samples were selected and downsampled to rates as low as 1 SNP per 10kb. Downsampling was repeated 100 times and then the resulting variant files were used in *MeiCOfi*. Both precision and sensitivity were computed by comparing the COs against the original list using bedtools.

### Testing *MeiCOfi*

We tested *MeiCOfi* on several different species with varying ploidy levels, for example, *Arabidopsis suecica*^3*2*^, *Arabidopsis thaliana*^33^, *Hordeum vulgare* L.^34^, *Oryza sativa*^13^, and *Solanum lycopersicum*^*30*^. *MeiCOfi* was also run on hyper-recombinant mutants, *zyp1 recq4ab*, in *Arabidopsis thaliana*^35^, and non-recombinant tetraploid hybrid *S. lycopersicum*^36^. More detailed information on sequencing data and reference genomes is available in Supp. Table. 1.

## Results

### Testing the pipeline

The *MeiCOfi* pipeline processes an individual sample to detect meiotic crossovers (Figure 1A). Combining multiple samples from the same population will generate a recombination landscape per chromosome. As shown in Figure 1B, an F1 hybrid shows a horizontal line, indicating genome-wide heterozygosity (e.g 0.5 for an F1 diploid hybrid), while an F2 offspring shows deviations from the expected ratio. The allele frequency patterns may be quite complex depending on the species of interest, some showing abrupt shifts due to the absence of markers or closely-spaced shifts due to double crossovers (Supp. Fig. 1). The basepair resolution (Figure 1C) of crossover detected by the tool enables comparison of COs with smaller features of the genomes, such as promoter regions or sequence motifs. Tested on simulated diploid samples with varying sequencing coverage (Figure 1D), the minimum F1-score is above 0.97, demonstrating the reliability of the tool. *MeiCOfi* presented an average precision score of 1 and a sensitivity of 0.94 at 0.5x coverage.

**Figure 1.**
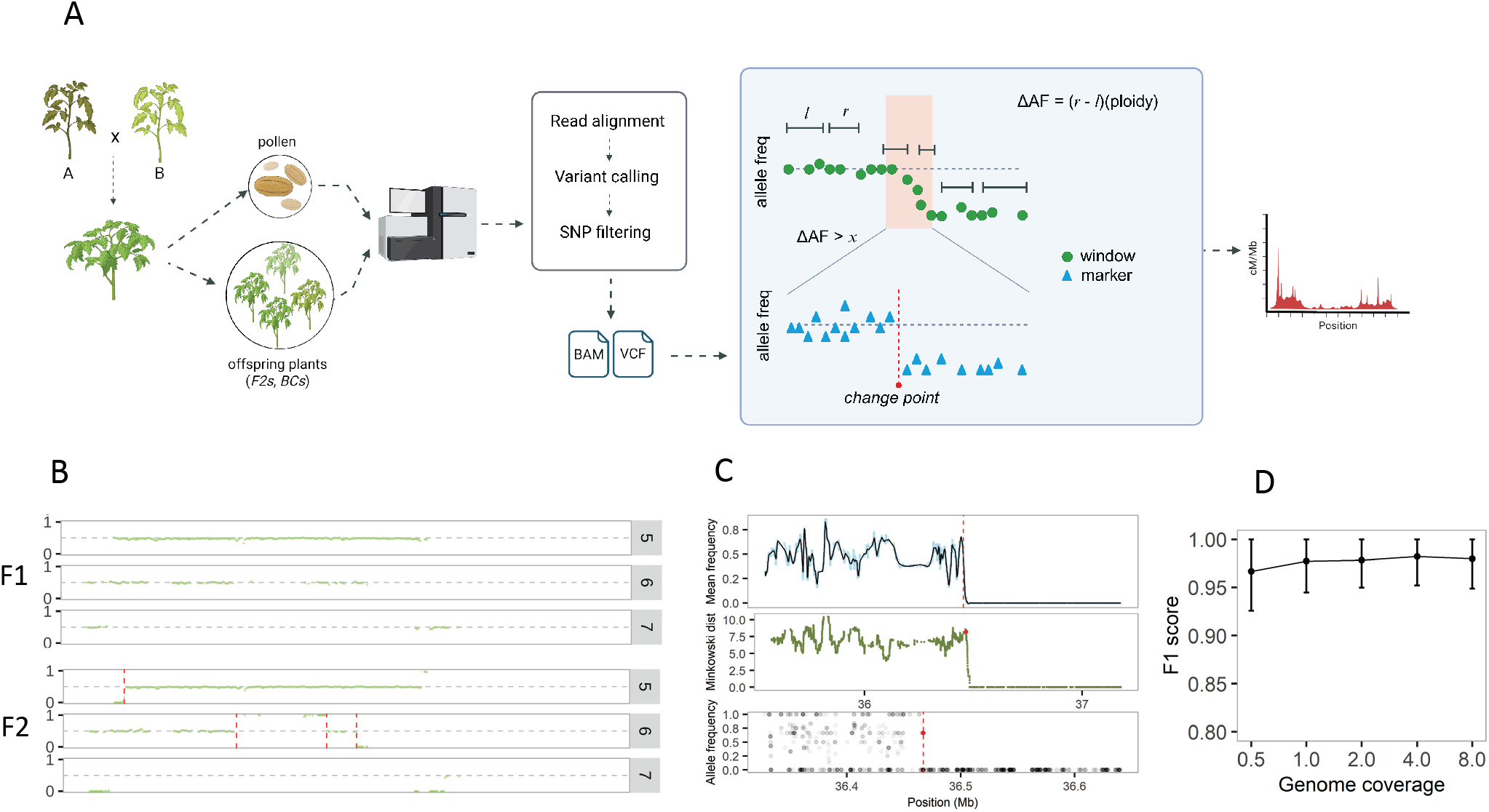
Detecting crossover events across genomes. A) MeiCOfi Pipeline. B) Allele frequency plot showing F1 and F2 samples, where each green dot represents the average frequency in a 500-kb window. Deviation from the frequency of 0.5 (heterozygous) indicates crossover (red line). Gap in chromosome 7 implies the absence of markers. C) The zoomed-in location of a crossover with raw allele frequencies. The three rows show the mean frequency per 50-SNP window, *Minkowski* distance between two consecutive non-overlapping 50-SNP window, and the raw allele frequency per SNP (black dots). The red line per row indicates the putative CO region, local distance maxima, and final crossover location, respectively. D) Performance in simulated diploid hybrids of *S. lycopersicum* cv. Mb-TMV x *S. pennellii* (Mb-Pen) with varying sequencing coverage.

To test on real data, we ran *MeiCOfi* on 95 recombinant samples of F2 tomato plants derived from a cross between *Solanum lycopersicum* cv. Moneyberg-TMV (MbTMV) and *Solanum pennellii* (LA0716). On average, these tomato samples have 3.3x coverage and 5 *MeiCOfi*-filtered SNPs per kilobase (Figure 2A). Shown in Figure 2B, the crossovers in different samples are precisely identified. There a total of 1,996 COs detected from 95 samples, with median resolution of 269 bp (Figure 2C). The resulting recombination landscape (Figure 2B) is consistent with previous data on *S. lycopersicum* x *S. pennellii* cross using offspring sequencing and genetic map^10,30,37^, accurately mapping recombination coldspots in distal chromosome ends. These coldspots have been associated with cross-specific features like inversions^30,37^. As reported previously^11,17,30,37^, most recombination in tomato is located at the distal ends of the chromosomes, while the pericentromeric regions have a significantly lower recombination rate. Crossover detection in sequencing data relies on genetic markers. Extremely low density of markers complicates the detection of recombination, limiting most recombination studies on hybrids with sufficient parental genetic differences. Using F2 tomato samples, we tested how the performance varies with reduced SNP density and the same parameter settings (Figure 2D). Any two consecutive COs located in a genomic region with no or an extremely low number of markers (e.g. pericentromeres) may not be detected. *MeiCOfi* may be fine-tuned by adjusting the parameters in species where markers are sparsely distributed across the genome.

**Figure 2.**
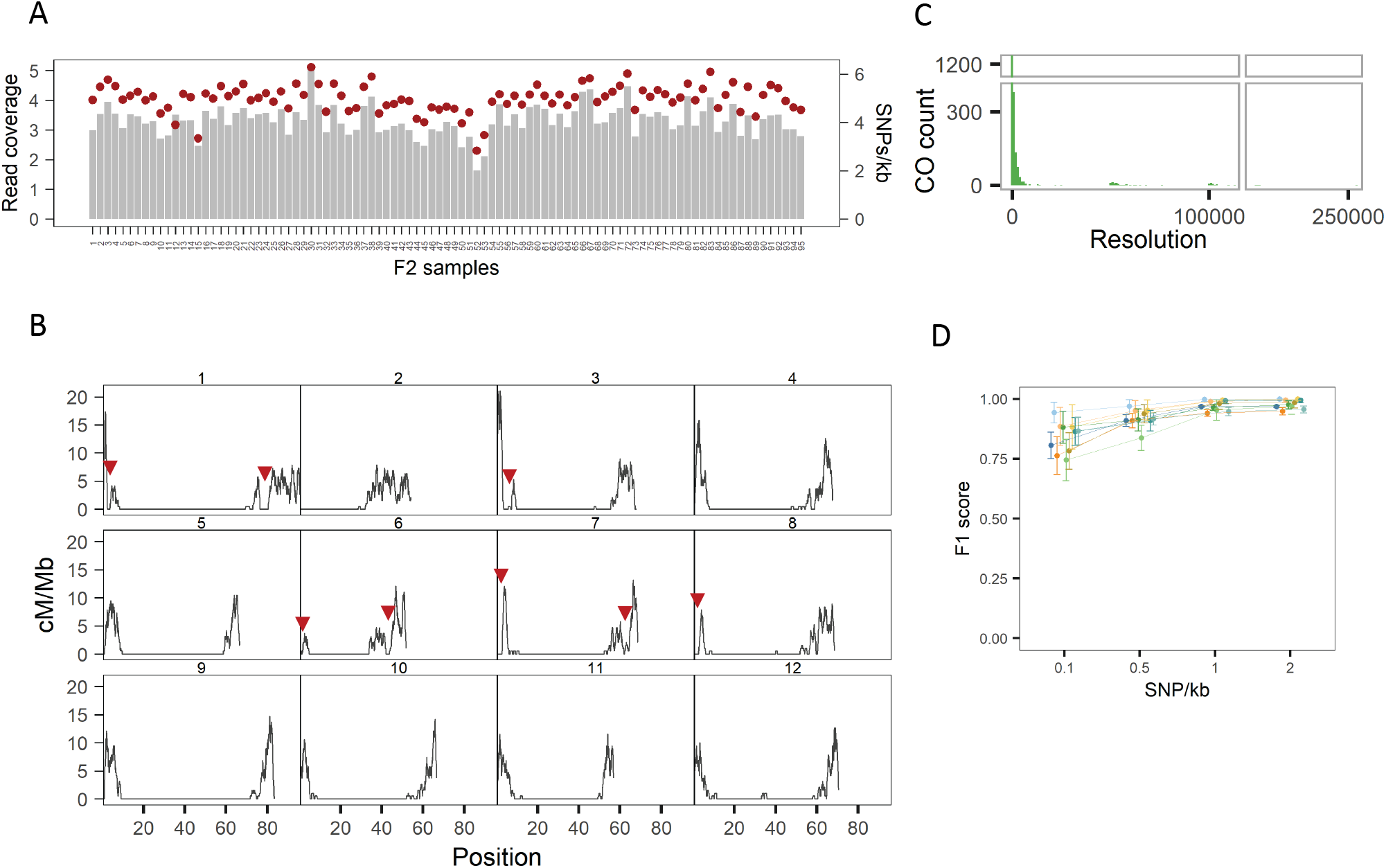
Recombination in interspecific tomato hybrids. A) Read coverage (gray bar) and density of scorable markers (red dots) in 95 F2 offspring of an interspecific hybrid cross between *S. lycopersicum* cv. Mb-TMV and *S. pennellii* (Mb-Pen). B) Recombination landscape of 95 F2 offspring. Red markers indicate the coldspots that match previously published recombination landscapes and genetic maps. C) Resolution of crossovers. Lower resolution in few COs is due to a low marker density in some genomic region. D) Performance of MeiCOfi in varying SNP density. Different density of SNPs is simulated 100 times in ten Mb-Pen diploid F2 samples.

### Detection in varying complexities

We ran *MeiCOfi* on offspring or gametes from different intra and inter-specific hybrids from diverse plant genera to test if it can identify recombination in variable genomic backgrounds and complexities. In Figure 3A, we show the SNP density and genome size of these hybrids. We sequenced intra-specific F2 offspring from a *S. lycopersicum* MbTMV x MicroTom hybrid, and F2 offspring from a commercial hybrid *Funtelle. MeiCOfi* detected COs in both hybrid genotypes but with lower sensitivity in Funtelle due to sparse distribution and low density of SNPs (Supp. Fig. 2). In interspecific hybrids between domesticated tomato and wild relatives, crossovers are detected accurately (Supp. Fig. 3). For recombination studies on closely-related parental genomes, it is crucial to ensure a sufficient number of markers and well-distributed markers in the genome.

**Figure 3.**
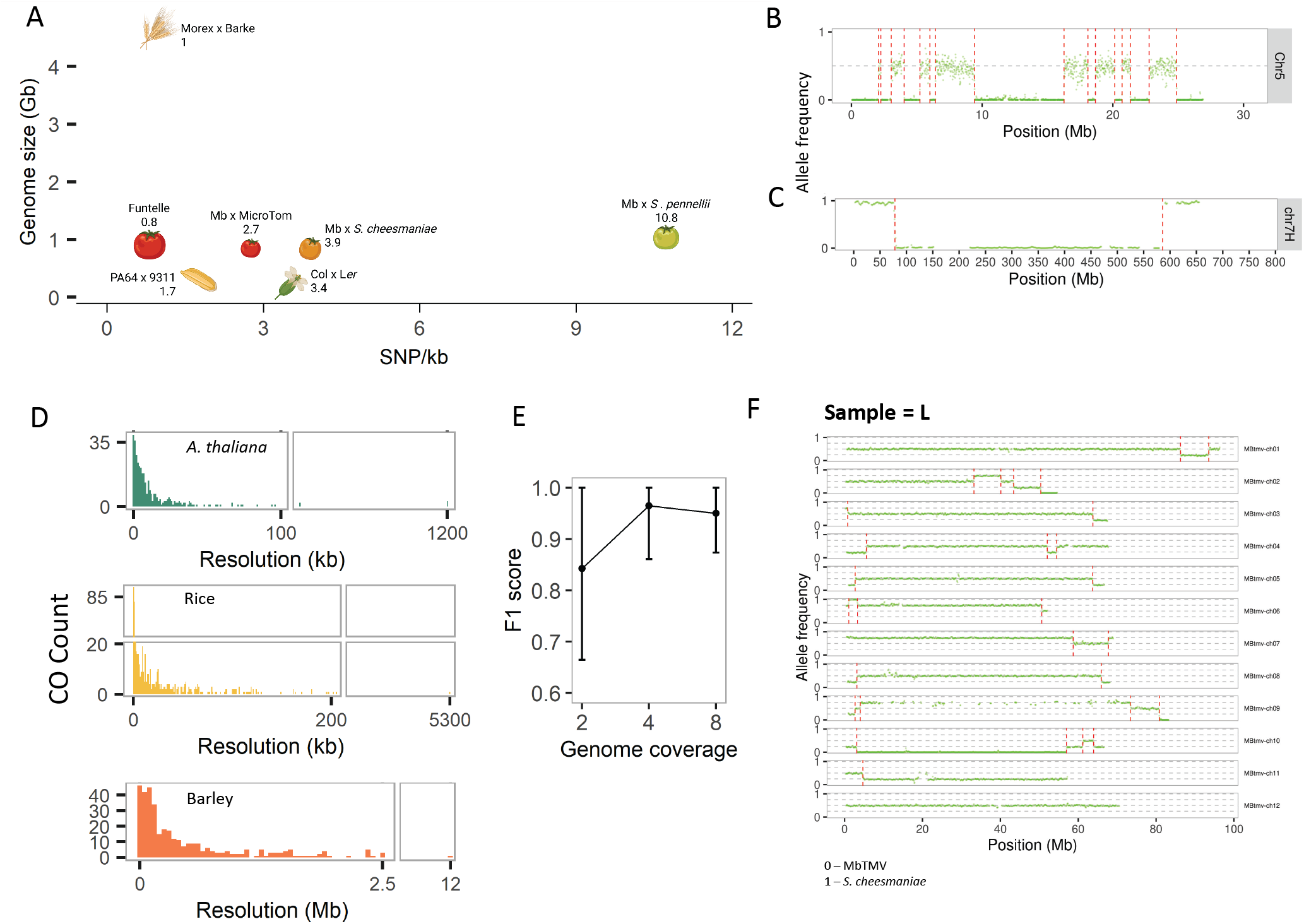
Applying MeiCOfi on different species and genome complexity. A) SNP density in hybrids of different species namely *Arabidopsis thaliana*, rice, barley and tomato (4 F1 hybrids of variable divergence). B) Hyper-recombination in a *zyp1 recq4a/b A. thaliana* Col-L*er* F1 hybrid backcrossed with *Col* plants (sample 5296_AB) with 16 COs (red line) in chromosome 5. C) Allele frequency plot generated from a haploid barley pollen from a Morex-Barke F1 hybrid (sample N4). Red line shows the crossovers. D) Resolution of crossover in 95 diploid Arabidopsis offspring (median=6kb), 24 diploid rice offspring (median=11kb), and 40 haploid barley pollen (median=233kb). E) Performance in simulated tetraploid tomato in *S. lycopersicum* cv. Mb-TMV x *S. pennellii* (Mb-Pen) background. F) Recombination in a synthetic tetraploid tomato made by doubling the genome of a *S. lycopersicum* x *S. cheesmaniae* F1 hybrid and selfing it.

We then tested *MeiCOfi* on *Arabidopsis thaliana*, specifically Col-0 x L*er* cross. Published sequence data on backcross plants^33^ were used to generate the recombination landscapes for male and female gametes (Supp. Fig. 4), indicating that our method can be used for other species. Furthermore, the tool was tested on rice^13^, which exhibits narrower low-recombining pericentromeres compared to tomato (Supp. Fig 5A & B). To further test the tool’s relevance on meiosis studies, we tested it on hyper-recombinant *zyp1 recq4a/b A. thaliana* Col-L*er* F1 hybrid samples with up to 12-fold more COs than wildtype^35^. Shown in a frequency plot, an Arabidopsis mutant chromosome exhibits an extremely high recombination rate (Figure 3B, Supp. Fig. 6A). *MeiCOfi* reported the precise boundaries of the two consecutive COs, which are less than 300kb apart (Supp. Fig. 6B).

Apart from analyzing diploid samples, we ran our method on 40 haploid pollen of barley (*Hordeum vulgare*), derived from a cross between cultivars *Morex* and Barke^34^. In contrast to diploids which have an expected allele frequency change of 0.5, in haploid samples the allele frequencies deviate from 0 to 1 (Figure 3C, Supp. Fig 7A). Similar to other crops, barley also have large pericentromeres with extremely low recombination rate (Supp. Fig. 7B). Compared to diploid samples, recombination in haploid gametes with 0.1x coverage can be accurately detected by *MeiCOfi* (Supp. Fig. 7C). A diploid sample contains homozygous and heterozygous markers while a haploid sample only classifies markers as alleles from either of the parent, allowing accurate marker genotyping even in low coverage. Furthermore, despite the lower number of SNPs (average = 0.26 SNP/kb) in barley due to low coverage (Supp. Fig. 7C), they are more spread out across the genome compared to Funtelle hybrid tomato, enabling better detection of COs. Compared with the megabase CO resolution based on a 5-Mb sliding window approach^34^, our method substantially improved the resolution of crossovers when markers are available (Figure 3D). A third of the COs now have resolution below 100-kb and one CO even has 7-bp resolution. Moreover, with higher coverage and SNP density, we observed higher CO resolution in Arabidopsis and rice than barley. Our findings indicate that low-coverage single gamete sequencing, when combined with *MeiCOfi*, can serve as a more cost-effective method for detecting recombination in cases where high resolution is not required.

### Recombination in polyploids

Detecting recombination in polyploids is another feature we added to our method. We first tested *MeiCOfi* on simulated allotetraploid genomes (Figure 3E). The average F1-score is above 0.95 for genomes with sequencing coverage above 4x and modest performance in 2x, indicating that for higher ploidy samples, higher coverage is preferred. To test on real tetraploid samples, we doubled the genome of a diploid MbTMV x *S. cheesmaniae* hybrid plant and self-pollinated it. The expected frequency change in the tetraploid plant is 0.25, as shown in Figure 3F. We demonstrated that *MeiCOfi* can detect recombination in tetraploid F2 recombinant tomato plants (Supp. Fig. 8) and absence of recombination in non-recombinant tetraploid hybrid tomato (Supp. Fig. 9)^36^. Apart from tetraploid genomes, we detected COs in a synthetic hexaploid *Arabidopsis suecica* sample (Supp. Fig. 10)^32^, indicating high sensitivity and precision. Further optimization for even higher ploidy levels, particularly in computing allele frequencies, is ongoing. Apart from the detection of COs in natural polyploids, our method can serve as a useful tool for exploring the inheritance mode in synthetic polyploids or testing for clonal reproduction in plants where synthetic apomixis has been engineered.

## Summary

We presented a new tool that can robustly detect crossover events in different species with varying marker densities, genome size and structure, recombination rate, and ploidy. The high sensitivity of detecting crossovers in both intra- and interspecific hybrid crosses and in polyploid genomes implies that *MeiCOfi* will be a useful tool for meiosis research, building recombination maps, and engineering polyploid crops. Robustness in detecting COs in low-coverage gamete data also provides an option to reduce the cost of generating a recombination map and avoid the laborious process of growing hundreds of plants. With the ability to work on diverse contexts, the users can design a variety of recombinant genome inputs for *MeiCOfi*.

## Contributions

RRF developed and tested the tool. JF generated the synthetic tetraploids, developed recombinant populations, and performed visual validation of the crossovers. TS generated variant files for rice and *A. thaliana*. YW developed the recombinant population derived from intraspecific hybrid plants. CJU supervised the research. RRF wrote the manuscript with contributions from JF, TS, and CJU.

## Acknowledgements

*S. cheesmaniae* LA1039 and *S. pennellii* LA0716 seeds were obtained from TGRC (UC Davis, California, USA). Moneyberg-TMV seeds were kindly provided for research purposes by Cilia Lelivelt and Maarten Verlaan (Rijk Zwaan, Netherlands). Funtelle F1 tomato seeds were kindly provided for research purposes by A. Voss (Syngenta, Germany). The authors thank Bruno Huettel for generating sequencing libraries and Raphael Mercier for providing insights on improving the tool.

## Data availability

The sequencing data for the diploid F2 tomato plants (PRJEB112205) and synthetic tetraploid F2 tomato plants (PRJEB112206) are deposited in ENA.

