## Supplementary Figures for "MeiCOfi: Meiotic CrossOver Finder in haploid, diploid, polyploid and hyper-recombinant genomes"

**Supplementary Figure 1. Types of allele frequencies shift**. Several types of allele frequency shifts are shown. Detecting crossovers at allele frequency shifts may be affected by SNP density and distribution, repeat content and genome complexity and ploidy. MeiCOFi parameters and sequencing coverage can be modified to overcome issues with crossover detection.

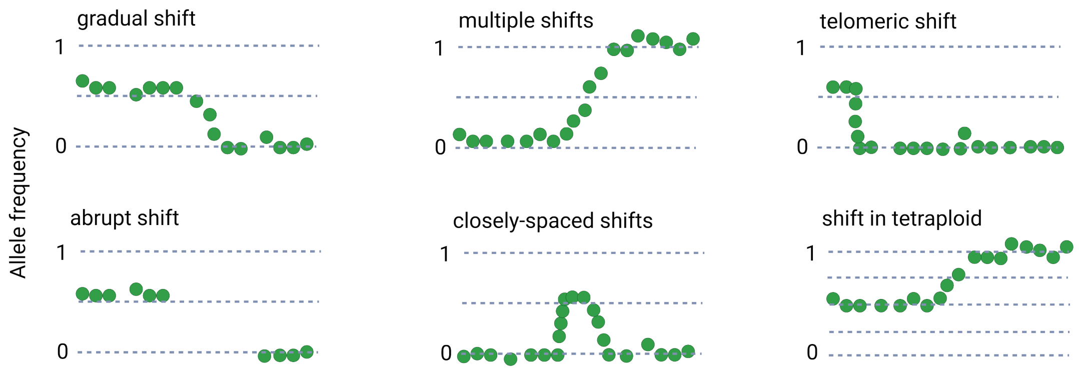

**Supplementary Figure 2. Crossover detection in intraspecific tomato hybrid F2s.** Allele frequency plots for (A) Three commercial hybrid *Funtelle* F2s (B) Six intraspecific tomato hybrid MbTMV x MicroTom F2s. The density of SNPs in Funtelle and MbTMV x MicroTom hybrids is 0.8 and 2.7 SNP/kb, respectively. In Funtelle hybrid, the number of SNP is low and SNPs are sparsely distributed along the chromosomes.

A

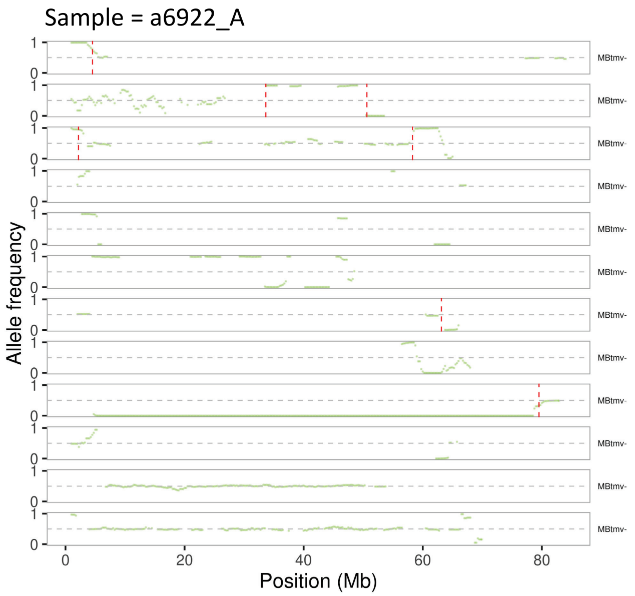


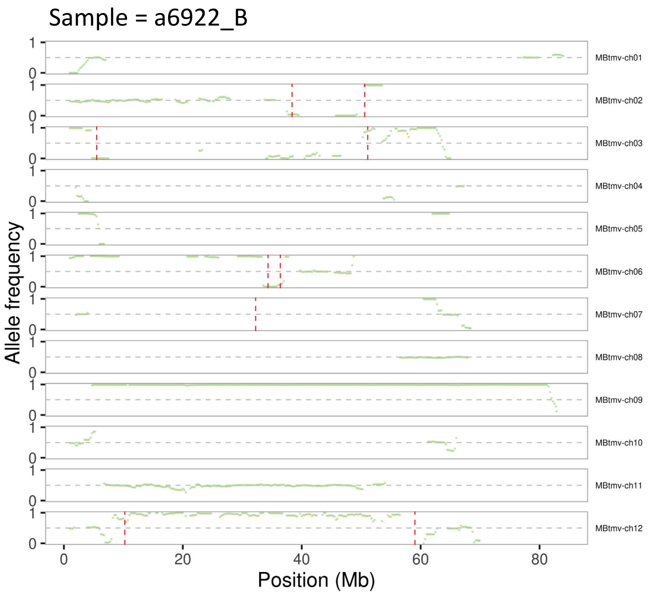


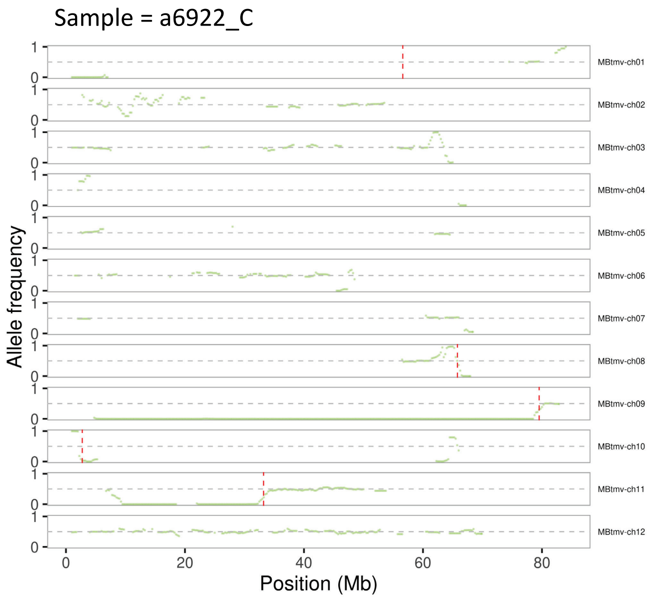

B

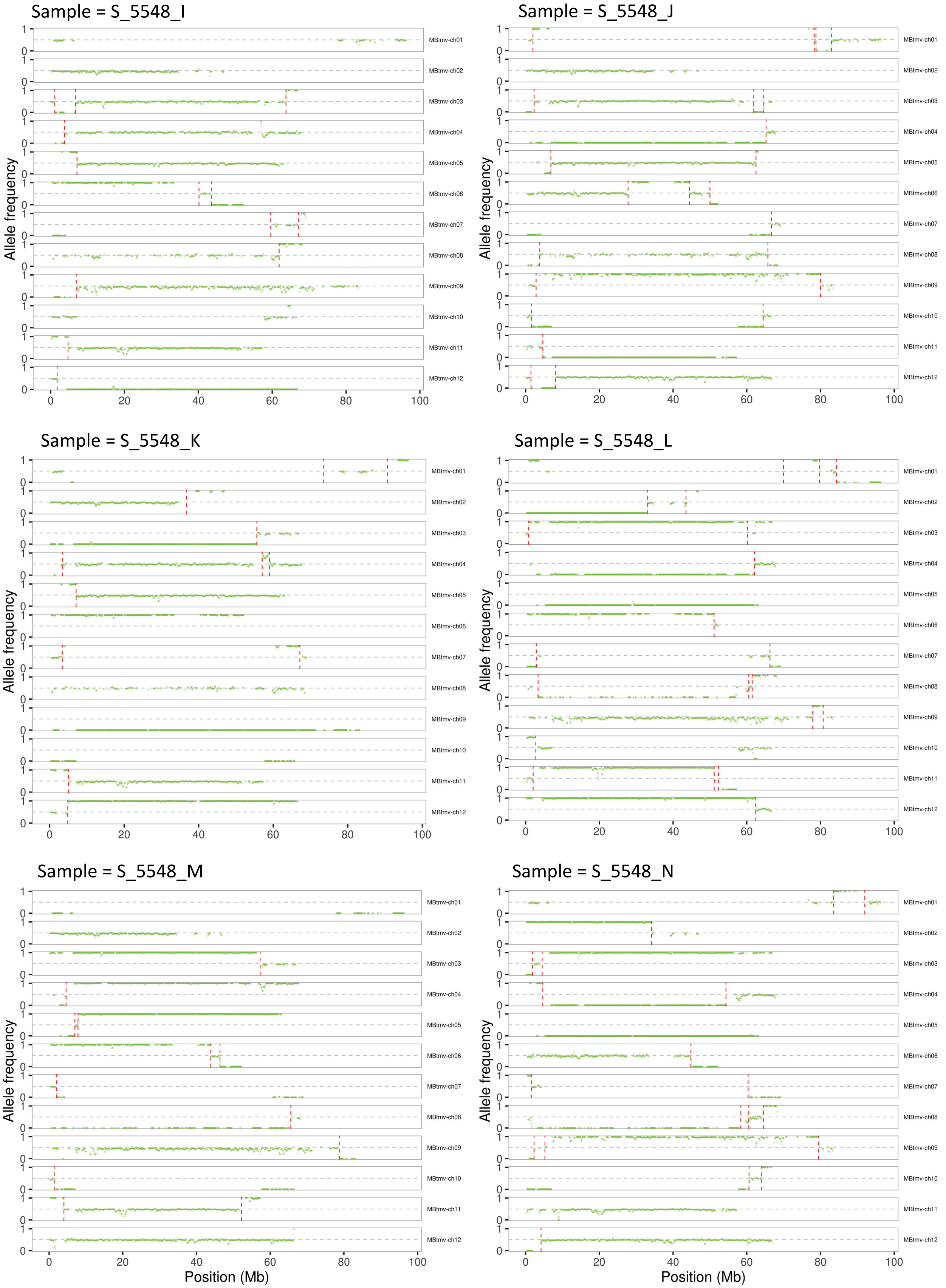
Sample = S_5548_I
Sample = S_5548_J
Sample = S_5548_K
Sample = S_5548_L
Sample = S_5548_M
Sample = S_5548_N
Allele frequency
Position (Mb)
0
1
0
20
40
60
80
100
MBtmv-ch01
MBtmv-ch02
MBtmv-ch03
MBtmv-ch04
MBtmv-ch05
MBtmv-ch06
MBtmv-ch07
MBtmv-ch08
MBtmv-ch09
MBtmv-ch10
MBtmv-ch11
MBtmv-ch12

**Supplementary Figure 3. Crossover detection in interspecific tomato hybrid.** Allele frequency plots for a (A) the first backcross generation of a *S. lycopersicum* cv. Moneyberg-TMV (MbTMV) x *S. cheesmaniae* (LA1039) hybrid backcrossed to MbTMV and (B) an F2 offspring of a hybrid of MbTMV and *S. pennellii* (LA0716).

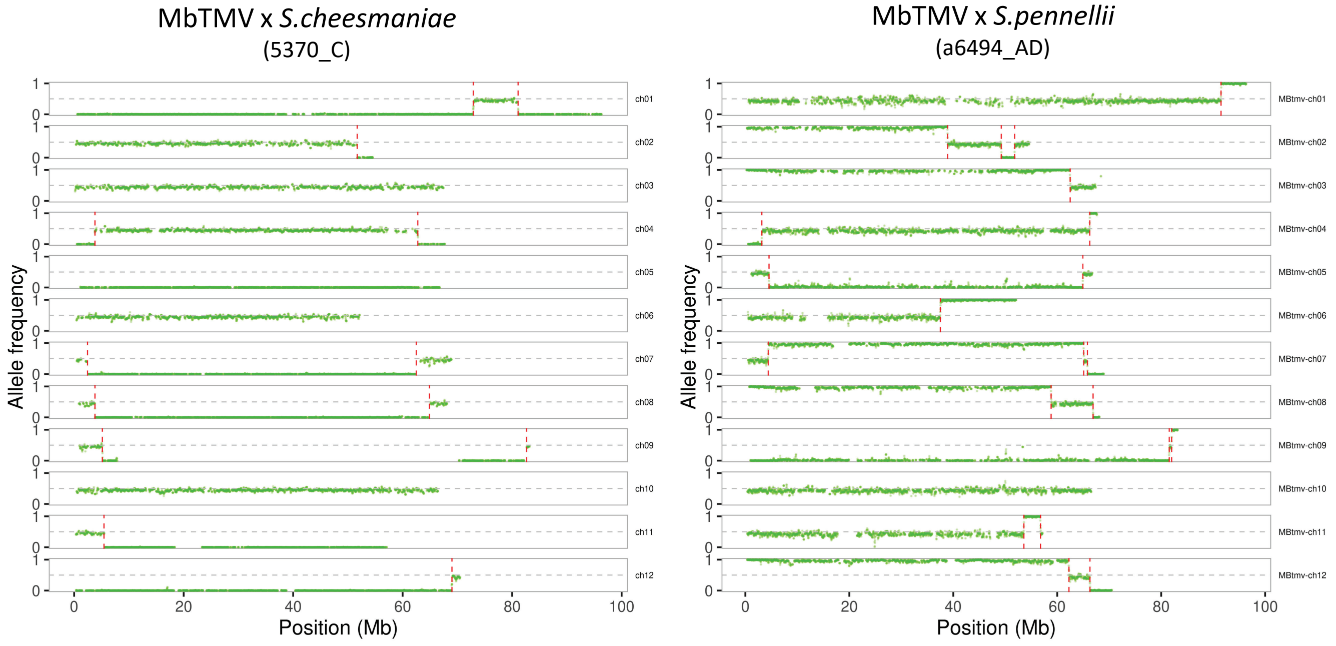

**Supplementary Figure 4. Recombination landscape in *A. thaliana***. Crossovers were detected in 47 male and 48 female *Arabidopsis* (Col-0 x *Ler*) backcross plants and used to generate a crossover landscape.

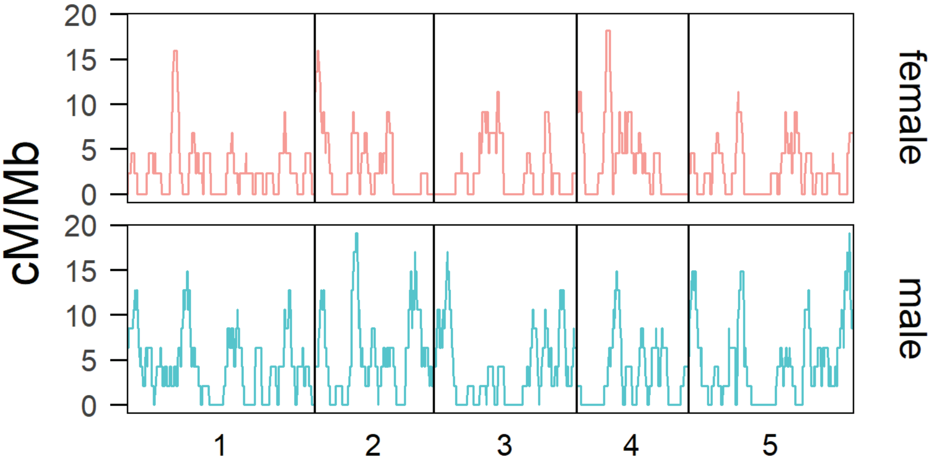

**Supplementary Figure 5. Recombination in rice genomes.** A) Allele frequency plot and crossovers in sample SRR1060368 (PA64 x 9311 hybrid). B) Recombination landscape derived from 24 rice F2 plants.

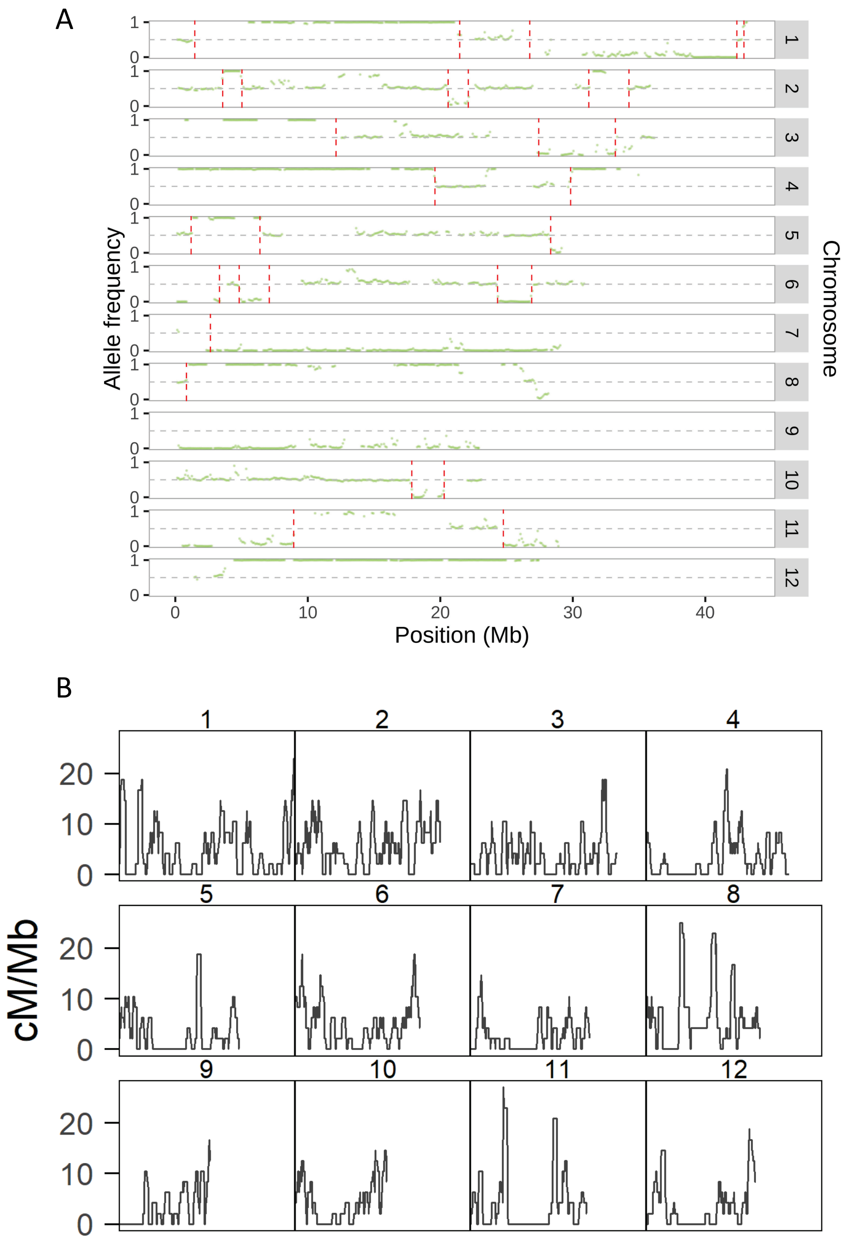

**Supplementary Figure 6. Hyperrecombination in *A. thaliana*.** A) Elevated recombination in *zyp1 recq4a/b A. thaliana* Col-*Ler* F1 hybrid backcrossed to *Col* plant where mutant sample 5296_AB and 5296_A have respective totals of 62 and 45 crossovers (red line). B) The zoomed-in graphics of the closely-spaced 2nd and 3rd crossovers in 5296_A chromosome 1. The three rows show the mean frequency per 50-SNP window, *Minkowski* distance between two consecutive non-overlapping 50-SNP window, and the raw allele frequency per SNP (black dot). The red line per row indicates the putative CO region, local distance maxima, and final CO location, respectively. (Parameters: window size = 30kb, step size = 15kb, minimum window = 6, number of SNPs = 20).

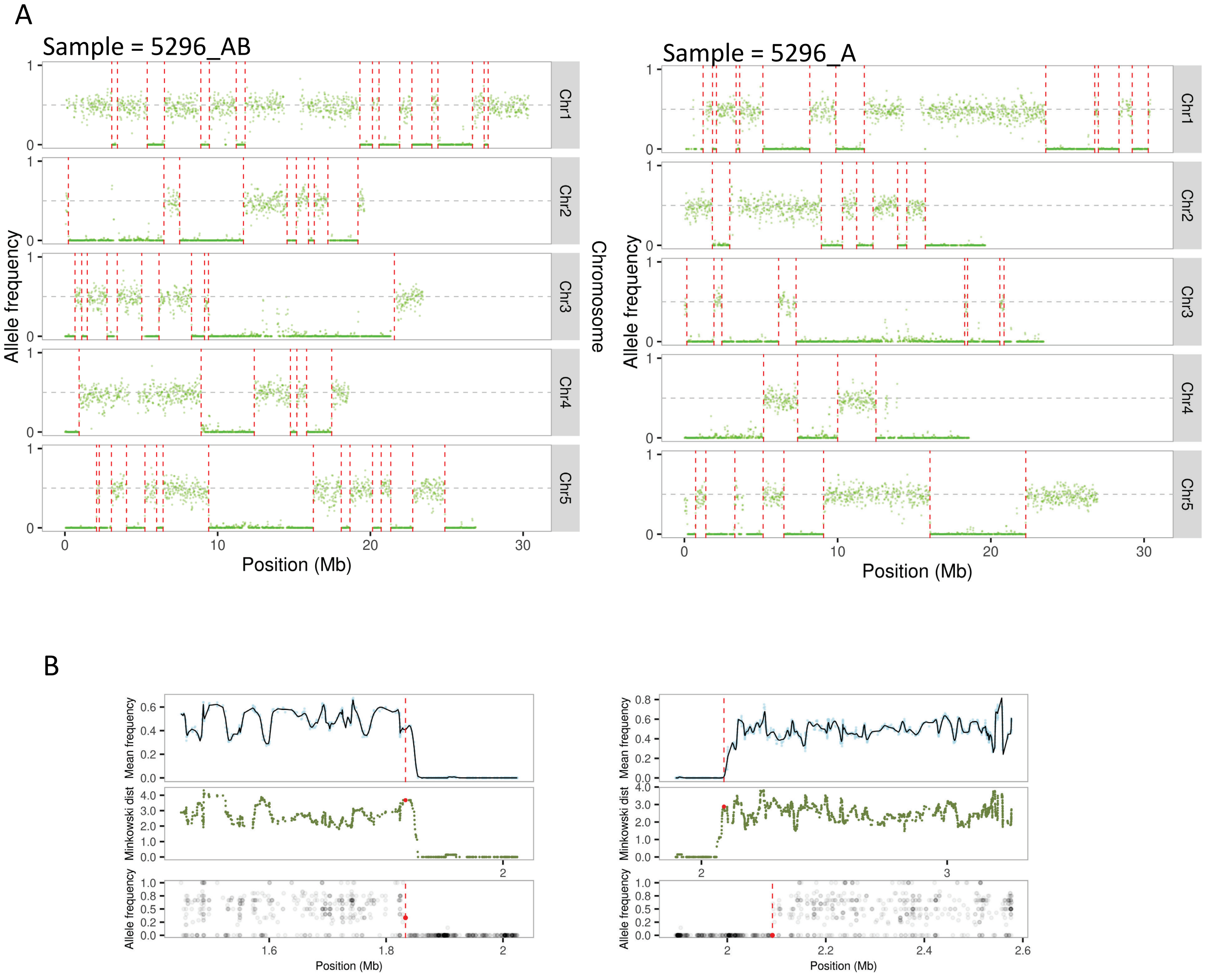

**Supplementary Figure 7. Recombination using barley pollen.** A) Allele frequency plot and crossovers in pollen sample N4 of *Hordeum vulgare* Morex x Barke F1 hybrid. B) Recombination landscape generated from 40 haploid pollen. C) Read coverage (gray bar) and density of scorable markers (red dots) in the pollen samples.

A

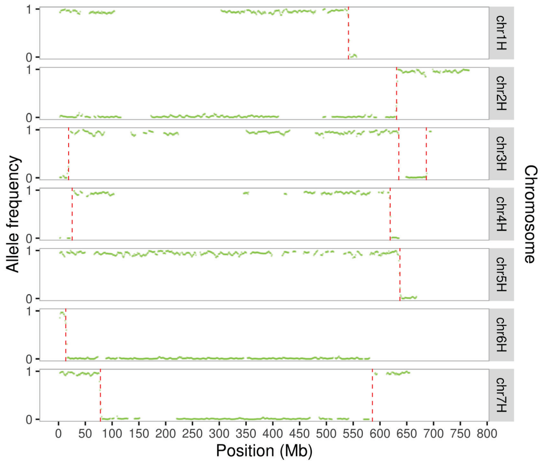


B

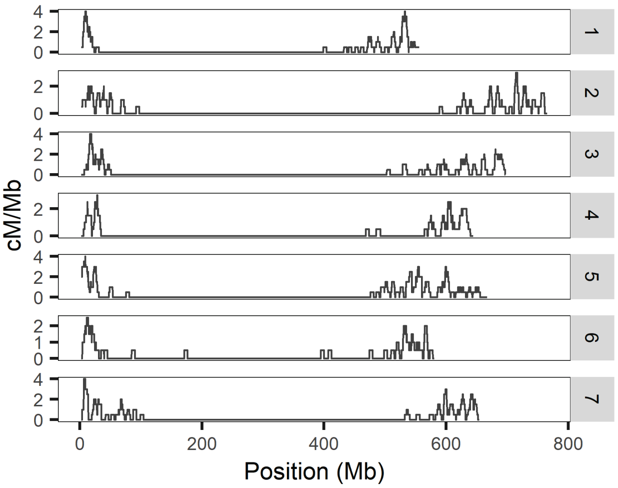


C

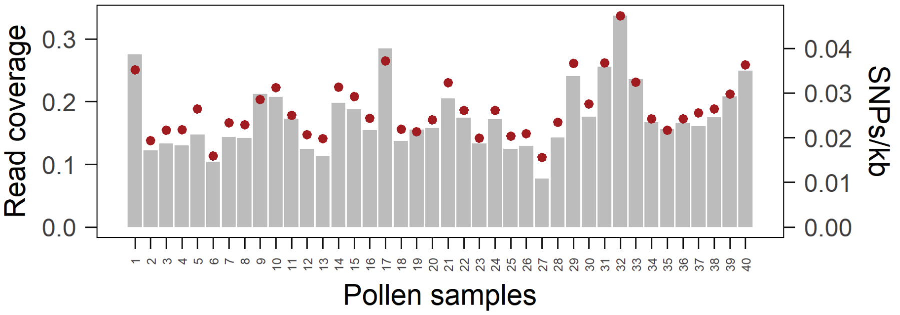

**Supplementary Figure 8. Crossovers in the genome of 13 synthetic hybrid tetraploid tomato F2 samples.** A 4x *Solanum lycopersicum* x *Solanum cheesmaniae* hybrid was self-pollinated and offspring were sequenced with read coverage between 7.5x and 21.6x. (Parameters: window size = 150kb, step size = 50kb, number of SNPs = 70).

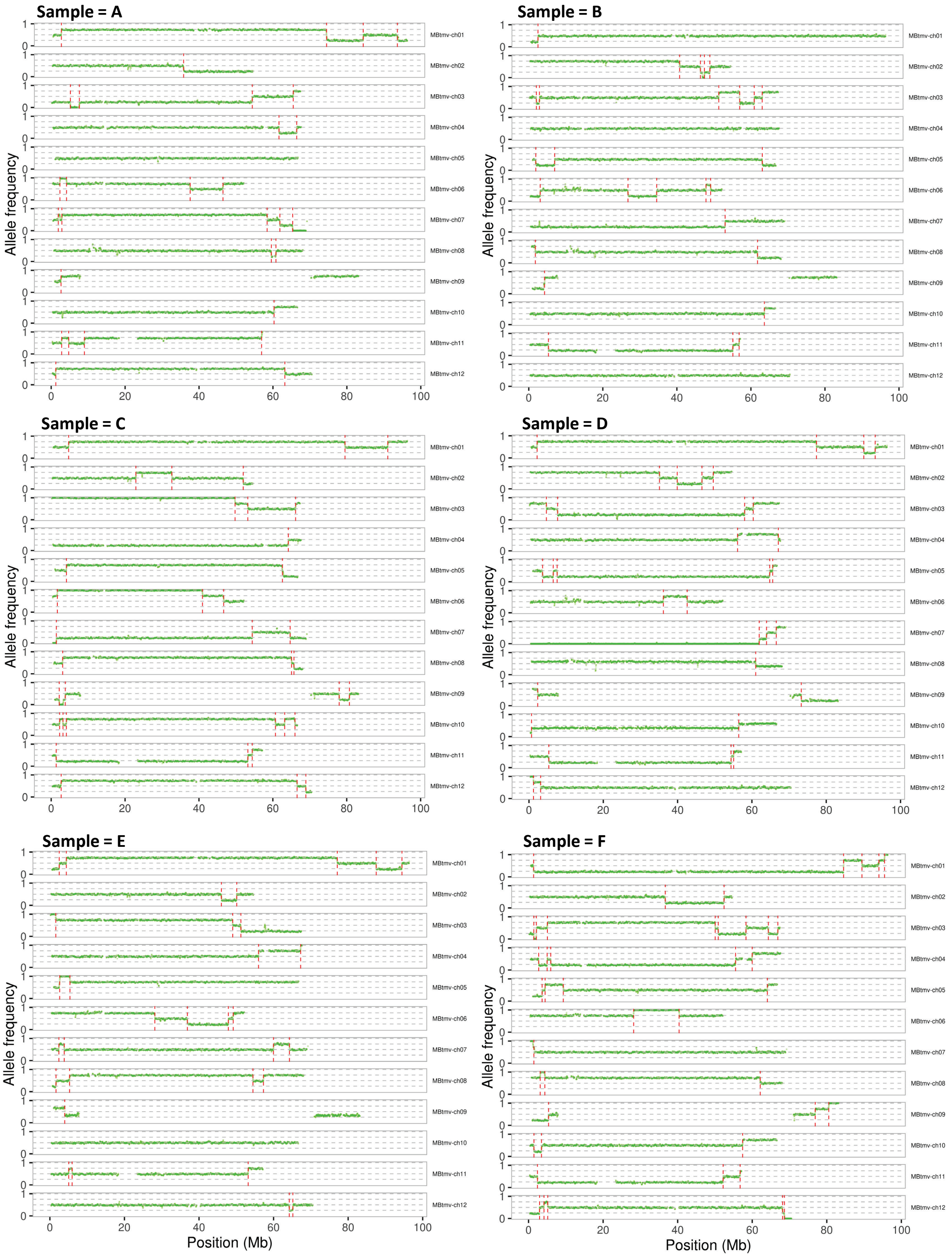

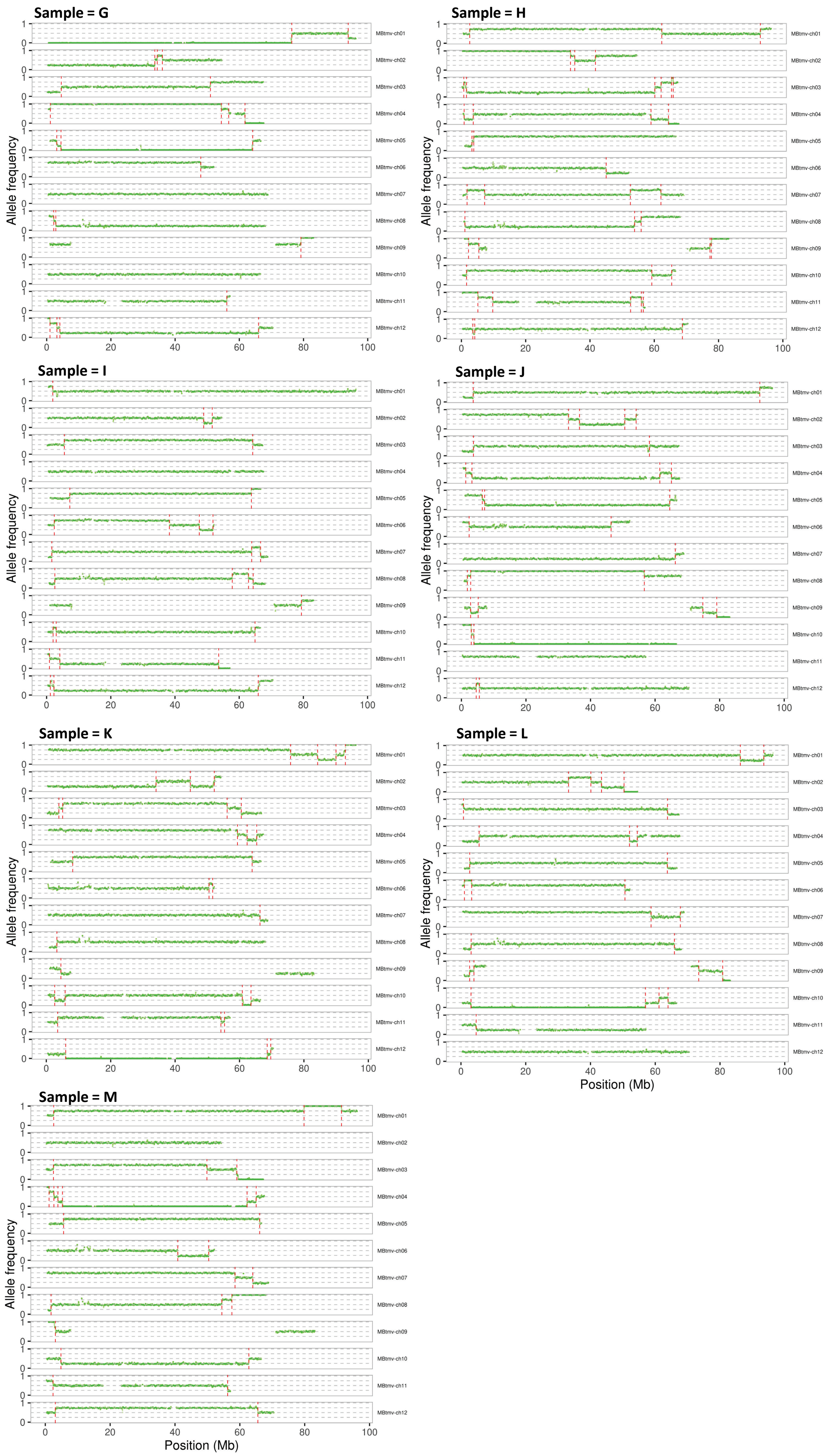

Sample = G
Sample = H
Sample = I
Sample = J
Sample = K
Sample = L
Sample = M
Allele frequency
Position (Mb)
MBtmv-ch01
MBtmv-ch02
MBtmv-ch03
MBtmv-ch04
MBtmv-ch05
MBtmv-ch06
MBtmv-ch07
MBtmv-ch08
MBtmv-ch09
MBtmv-ch10
MBtmv-ch11
MBtmv-ch12
0
1
20
40
60
80
100

**Supplementary Figure 9. Absence of crossovers in tetraploid tomato *MiMe* offspring samples**. *Solanum lycopersicum* cv. MbTMV x *Solanum lycopersicum* cv. MicroTom diploid plants were engineered to undergo *Meiosis instead of Meiosis.* Tetraploid offspring were produced by the fertilization of diploid clonal eggs by diploid clonal sperm cells. MeiCOfi confirmed the absence of crossovers in the samples. (Parameters: window size = 5Mb, step size = 100kb, number of SNPs = 20)

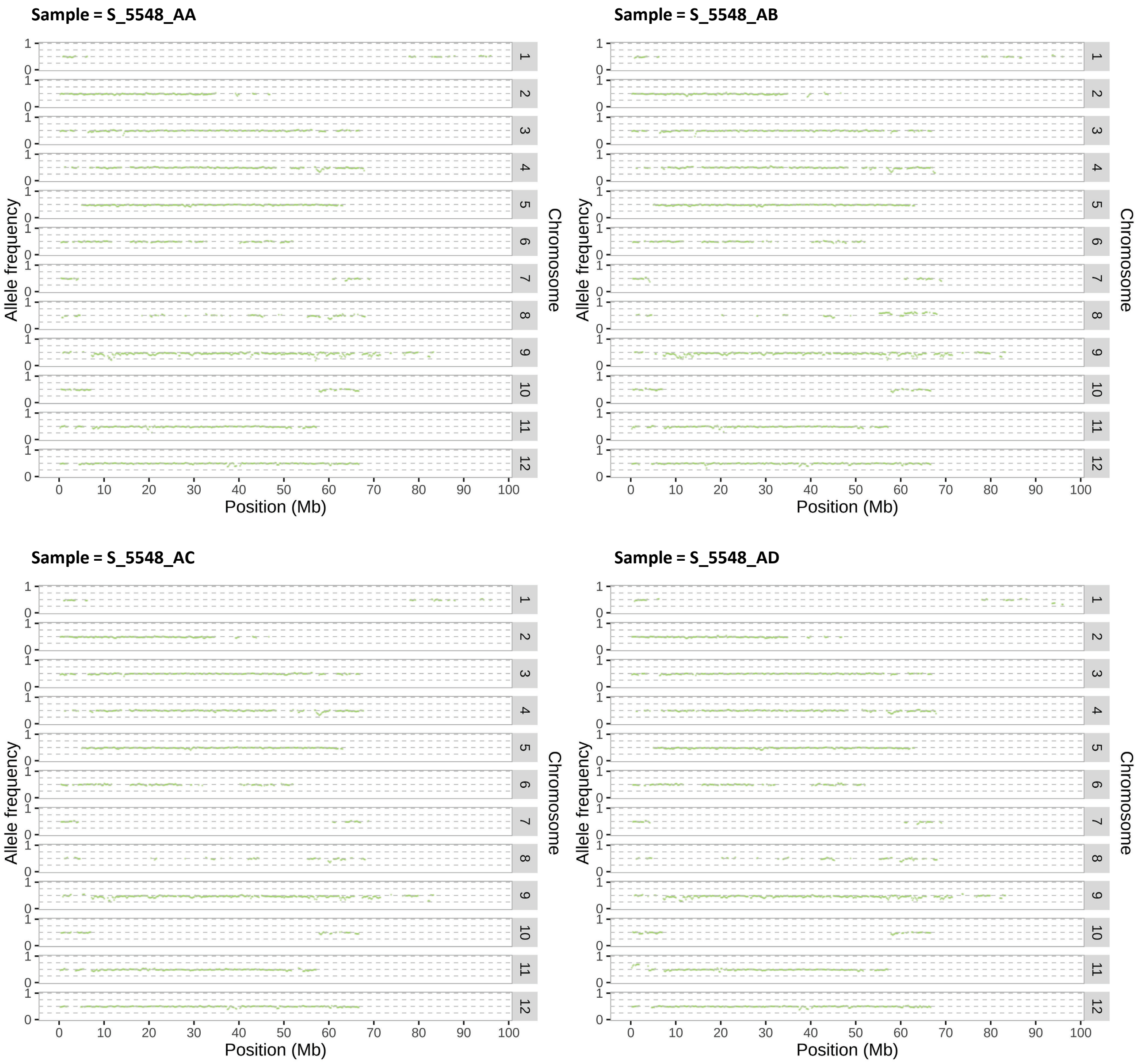

**Supplementary Figure 10. Crossovers in the hexaploid genome of an F2 offspring of a synthetic *Arabidopsis suecica* lineage**. A hybrid between A. thaliana (2x) x A. arenosa (4x) underwent spontaneous genome duplication, producing a hexaploid plant. (Parameters: window size = 500kb, step size = 50kb, number of SNPs = 30).

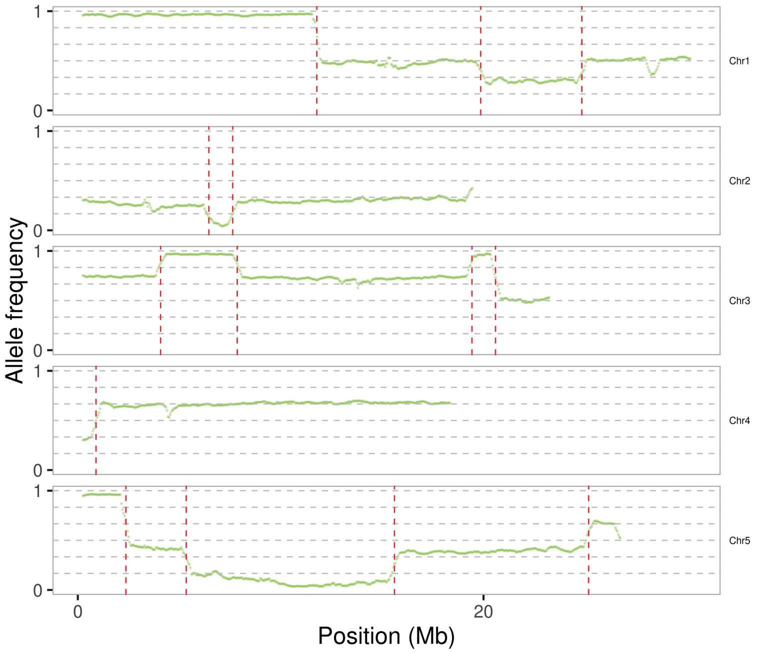


0 – *A. thaliana*
1 – *A. arenosa*
